# Automated protein-protein structure prediction of the T cell receptor-peptide major histocompatibility complex

**DOI:** 10.1101/2022.06.01.494331

**Authors:** Zachary A. Rollins, Matthew B. Curtis, Roland Faller, Steven C. George

## Abstract

T Cell Receptor (TCR) recognition of a peptide-major histocompatibility complex (pMHC) is a crucial component of the adaptive immune response. The identification of therapeutically relevant TCR-pMHC pairs is a significant bottleneck in the implementation of TCR-based immunotherapies but may be augmented by computational methodologies. The ability to computationally design TCRs to target a specific pMHC will require an automated integration of next-generation sequencing, protein-protein structure prediction, molecular dynamics (MD), and TCR ranking. We present a generic pipeline to evaluate patient-specific, sequence-based TCRs to a target pMHC. Using the three most frequently expressed TCRs from 16 colorectal cancer patients, we predicted the protein-protein structure of the TCRs to the target CEA peptide-MHC using Modeller and ColabFold. Then, these TCR-pMHC structures were compared by performing an automated molecular dynamics equilibration. ColabFold generates starting configurations that require, on average, a ~2.5X reduction in simulation time to equilibrate TCR-pMHC structures compared to Modeller. In addition, the structural differences between Modeller and ColabFold are demonstrated by an increase in root mean square deviation (~0.20 nm) between clusters of equilibrated configurations, which can impact the number of hydrogen bonds and Lennard-Jones contacts between the TCR and pMHC. Finally, we identify a TCR ranking criteria that may be used to prioritize TCRs for evaluation of *in vitro* immunogenicity.

## INTRODUCTION

Cytotoxic (CD8+) T cells are part of the adaptive immune system and eradicate potentially harmful cells – including cancer cells – by recognition of the peptide-major histocompatibility complex (pMHC) on target cells. Tumor specific pMHCs are comprised of a peptide derived from a mutated and/or aberrantly expressed intracellular protein (1) presented to the cell membrane in a pocket formed by the MHC ⍰ and β chains (2). The diversity of peptide-MHCs (~10^6-12^)(3) is matched by the diversity of TCRs (>10^20-61^) (4,5) through random V(D)J recombination of the hypervariable complementarity determining regions (CDRs). The function of the adaptive immune response ultimately depends on the ability to produce appropriate immunogenic TCRs (on-target) while minimizing response to self pMHCs (off-target effects).

Despite breakthrough clinical potential for TCR-T cell therapies in solid tumors (6–10), the implementation is hindered by three central challenges: 1) identifying tumorspecific pMHC ligands; 2) matching immunogenic TCRs with identified pMHCs, and 3) minimizing off-target effects (11). Combining next generation sequencing and machine learning, significant advancements have been made to identify and rank tumor-specific pMHC ligands(12–14), thus addressing the first challenge.

Addressing the second challenge has been difficult as the identification of patientspecific TCR repertoires has involved methods that are low-throughput or limited to a single chain (15–17). However, recent breakthroughs in single-cell sequencing allow determination of the CDR3 regions of the ⍰ and β chain of the TCR in a high-throughput manner (18–21). This technological breakthrough facilitates an unprecedented exploration of the vast TCR information space and allows the scientific community to refocus attention on fundamental questions related to recombination, maturation, and intersecting diversity of patient-specific TCR repertoires. This advance also provides an opportunity to leverage machine learning to predict TCR antigen binding specificity from primary amino acid sequence (22) or from structural features of TCR-pMHC homology models (23). However, the training sets to characterize and rank TCRs by their immunogenicity are restricted by either insufficient data on the relevant TCR-pMHC binding parameters (24–30) or a limited number of known TCR-pMHC structures (~400 on STCRab) (31).

Despite significant advances in protein-protein structure prediction (32–38), the prediction of TCRs bound to a target pMHC from patient-specific sequences is fundamentally biased to the features of known protein structures. Moreover, ranking TCRs is not possible without detailed information on the relationship between bond strength and immunogenic response (24–30). Previously, we have identified several physiochemical parameters of the TCR-pMHC interaction that correspond with immunogenicity (28,39). Herein, we generate an automated pipeline to assess TCRs to a target pMHC (**Figure 1**). This pipeline begins with single-cell sequencing to identify the amino acids in the CDR3⍰β loops from T cells resected from the tumors of 16 colorectal cancer (CRC) patients(21). Next, we restrict the carcinoembryonic (CEA) peptide (CEA571-579:YLSGANLNL) to the MHC (HLA-0201), known to be expressed in CRC patients (11,40–42) and predict several TCR-pMHC complexes using TCRs sequenced from patients (21). The predicted protein structures are equilibrated at physiological conditions by molecular dynamics simulations in Gromacs (43,44) and an automated equilibration (45) is implemented to assess the starting structures from either Modeller (35,36) or the recently developed ColabFold (32–34) (**Figure 1**). Our results demonstrate that ColabFold results in structures that are ~2.5X faster to equilibrate, and thus may reduce computational cost compared to Modeller. However, the clusters of structures generated by Modeller and ColabFold are consistently divergent despite structural equilibrium. Moreover, we provide potential criteria for ranking the TCRs after structural equilibration including the number of hydrogen bonds and Lennard-Jones contacts. This methodology is generally applicable to identify TCRs with relevant and quantifiable binding parameters to a target pMHC.

**Figure 1:**
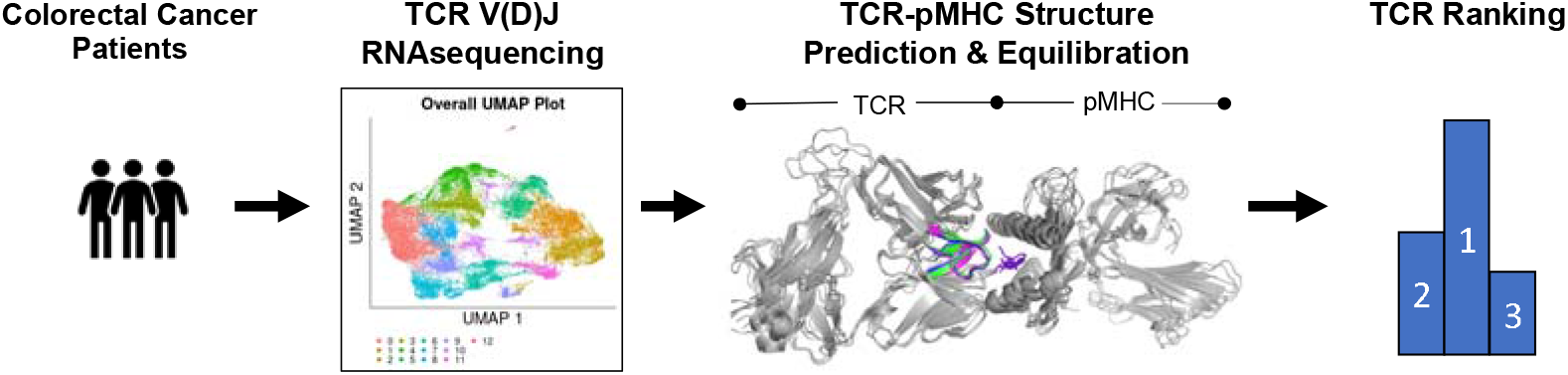
Process flow diagram for the protein-protein structure prediction of TCRs to a target pMHC. The process begins with single cell V(D)J RNA sequencing of the T cells from resected tumors of 16 colorectal cancer patients (**left**). Then, protein-protein structure prediction of TCRs sequenced from patients bound to a pMHC (HLA-A2) with a restricted target peptide CEA_571-579_ (**middle right**) is performed. Finally, we run molecular dynamics simulations to equilibrate the structure and rank TCRs based on the number of interactions at equilibrium (**right**).

## METHODS

### Single-cell RNA V(D)J sequencing of CRC patient T cells

T lymphocyte single cell RNA-Seq data was made available to us from the Han group and has been previously published (20,46). Raw data was first put through a quality control process to exclude cells with less than 200 unique genes, more than 7,500 unique genes, and/or more than 10% mitochondrial gene expression. Additionally, any genes that were present in fewer than 3 total cells were excluded from downstream analysis. All single-cell analysis was performed using the Seurat pipeline(47–50) in the R computing environment. T lymphocytes were clustered with UMAP(51) using 0.3 as the value for the “resolution” parameter, which yielded a total of 13 unique clusters. Cytotoxic T lymphocyte clusters were identified by expression of *CD3D* and *CD8A*, and the absence of *CD4* expression. TCR CDR3α and CDR3β sequences from the 10X Genomics 5’ VDJ analysis pipeline were then matched to their corresponding cells for downstream analysis.

### TCR-pMHC protein-protein structure prediction

The starting structures for TCR1, TCR2, and TCR3 were generated using Modeller v10.1 (35,36) and ColabFold v1.2.0 (32–34). The primary amino sequence used for multiple sequence alignment was derived from the DMF5 TCR bound to the HLA-A2 (MHC) restricted MART1 (PDB:3QDJ) (52). For sequence alignment, the CDR3⍰ (CAVNFGGGKLIF), CDR3β (CASSLSFGTEAFF), and MART1 peptide (AAGIGILTV) were substituted with the respective CDR3 loops found from patient TCR clonotypes (**Table 1**) and the CEA571-579 peptide (YLSGANLNL) known to be restricted to the HLA-A2 (MHC). For Modeller, the MART1 (PDB:3QDJ) crystal structure was used as the template, 10 model structures were generated from the alignment of the respective TCR, and the structure with lowest DOPE score (53) was selected for MD equilibration. ColabFold (34) is, in part, a server that performs rapid MSA/homology search combined with the trained network architecture of AlphaFold2 (32,33) for prediction of the 3D atomic coordinates of folded protein structures. For ColabFold, 5 model structures were generated from the alignment of the respective TCR, and the structure with the highest pTMscore (32,33) was selected for MD equilibration.

**Table 1:**
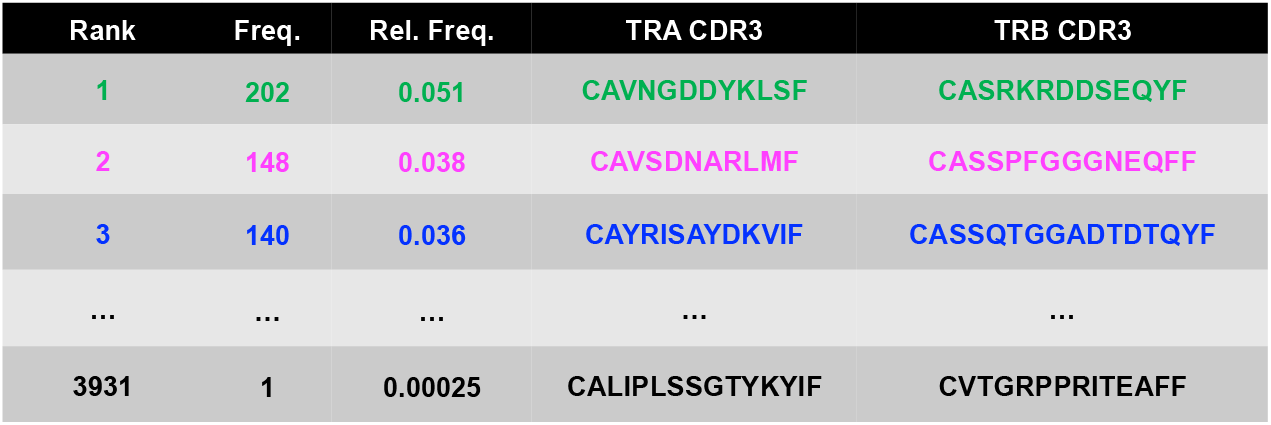
Rank order of CD8^+^ TCR⍰β clonotypes by relative frequency. This includes the frequency and relative frequency that the clonotype was identified as well as the CDR3⍰β primary amino acid sequence associated with the TCR⍰β clonotype.

### Molecular dynamics: setup, energy minimization, and equilibration

The predicted structures are then used as starting configurations for a 7 step molecular dynamics pipeline to determine their equilibrated structures at physiologic conditions. All MD simulations were performed with Gromacs 2019.1 (43,44) using the CHARMM22 with CMAP force field (54), orthorhombic periodic boundary conditions, and in full atomistic detail. (1) The residue protonation states are determined by calculating pKa values using propka3.1 (55,56) and deprotonated if pKa values are below physiologic 7.4 pH. (2) The properly protonated or deprotonated protein structures are solvated in rectangular water boxes large enough to satisfy minimum image convention using the TIP3P water model (57). (3) Na^+^ and Cl^-^ ions are added to reach physiologic salt concentration ~150mM and neutral charge. Box sizes were ~11.4 x 10.9 x 13.8 nm with ~50,000 water molecules, ~300 ions, and ~150,000 total atoms. Full specifications can be found in the Dryad repository: https://doi.org/10.25338/B83S70. (4) To avoid steric clashes, steepest descent energy minimization (emtol = 1000kJ/mol/nm) is performed. (5) To relax solute-solvent contacts, a 100 ps simulation is run in the constant volume ensemble (NVT) with 0.2 fs timestep (T = 310K). Temperature is maintained by coupling protein and nonprotein atoms to separate baths using a velocity rescale thermostat (58) with a 0.1 ps time constant. (6) To maintain pressure at 1.0 bar, a 100 ps simulation is run in the constant pressure (NPT) ensemble using isotropic Berendsen pressure coupling (59), a 2.0 ps time constant, and 2 fs timestep. Steps (5)-(6) used position restraints (harmonic force constant = 1000 kJ/mol/nm^2^) on all protein atoms. (7) Equilibration MD simulations were conducted for 150-300 ns with no restraints. Equilibration runs were extended in 50 ns increments until the root mean square deviation of the TCR-pMHC complex was in equilibrium for a minimum of 50 ns determined by the variance-bias trade-off algorithm (45). To maintain temperature and pressure during the equilbration runs, the Nose-Hoover thermostat (60) and Parrinello-Rahman barostat (61) were used with time constants 2.0 and 1.0 ps, respectively. The isothermal compressibility of water is used as 4.5×10^-5^ bar^-1^. Simulations used the Particle Ewald Mesh algorithm (62,63) for long-range electrostatic calculations with cubic interpolation and 0.12 nm grid spacing. Shortrange nonbonded interaction were cutoff at 1.2 nm. All water bond lengths were constrained with SETTLE (64) and all other bond lengths were constrained using the LINCS algorithm (65). The leap-frog algorithm was used for integrating equations of motion with a 2 fs time step.

### Data and Statistical Analysis

Selected TCR-pMHC structures from MD trajectories were visualized using the Pymol Molecular Graphics System v2.4.0 (Schrodinger, LLC; New York, NY). The selected frames for visualization were chosen to be from the top three clusters (2 from each cluster) after TCR-pMHC structure equilibration (**Figure S1-S2**). This resulted in a total of 12 structures for each TCR: 6 from Modeller (TCR1: Cluster 1, 2, & 7, TCR2: Cluster 1, 9, & 12, and TCR3: Cluster 1, 6, & 9) and 6 from ColabFold (TCR1: Cluster 1, 2, & 4, TCR2: Cluster 1, 3, & 4 and TCR3: Cluster 1, 3, & 6) (**Figure S3-S5**). The clusters were selected because they were after the simulation time required for equilibrium (e.g., for Modeller TCR1, cluster 1,2, & 7 were chosen because clusters 3-6 are only dominant before reaching equilibrium at 249 ns simulation time). The all-to-all alignment of TCR-pMHC structures was performed in Pymol using the align command on the C^⍰^ atoms to compute RMSD between pairs of structures. Data analysis from MD simulations was performed with tools from the Gromacs suite(43): gmx make_ndx, gmx hbond, gmx rms, gmx rmsf, and gmx cluster. These results were complemented with a secondary analysis utilizing python packages for data handling and visualization including: numpy(66), pandas(67), matplotlib(68), GromacsWrapper(69), scipy(70), and pingouin(71), and pymbar (45). Custom bash shell, python, and R scripts relevant to the production of figures are deposited in a Github repository: https://github.com/zrollins/TCR_homology.git. The geometry of a Lennard-Jones contact is defined as a distance of less than 0.35 nm between atoms. Results were presented as mean ± SEM. As indicated in figures, statistics were performed in python using scipy for one-way analysis of variance (ANOVA), and pingouin for pairwise Tukey-HSD post-hoc tests. Detailed outputs of statistical analysis were written to excel and are provided in a Dryad repository: https://doi.org/10.25338/B83S70.

## RESULTS

To design TCRs to target the CEA_571-579_ peptide restricted to the HLA-A2 (MHC), TCR clonotypes were identified utilizing the single cell RNA V(D)J sequenced T cells resected from colorectal tumors of 16 CRC patients (21). This technique identifies TCR clonotypes by matching the amino acids from the CDR3 regions of the ⍰ and β chain of the TCR (18–21). First, T cells were identified by the expression of *CD3D* and *CD8A* (and the absence of *CD4* expression), as only CD8*+* T cells can bind to the HLA-A2 (MHC). Of the 37,931 T cells analyzed (**Figure 2Ai**), there were 9,709 *CD3D+CD4-CD8A+* T cells (corresponding to clusters 2, 6, 7, 9, and 11; **Figure 2Aii-iv**), and corresponding TCR⍰β clonotypes (**Figure 2B, Table 1**). The three most frequently identified TCRs (**Table 1**) from patient tumors (denoted clonotype TCR1, TCR2, and TCR3) were then used to predict TCR-pMHC structures.

**Figure 2:**
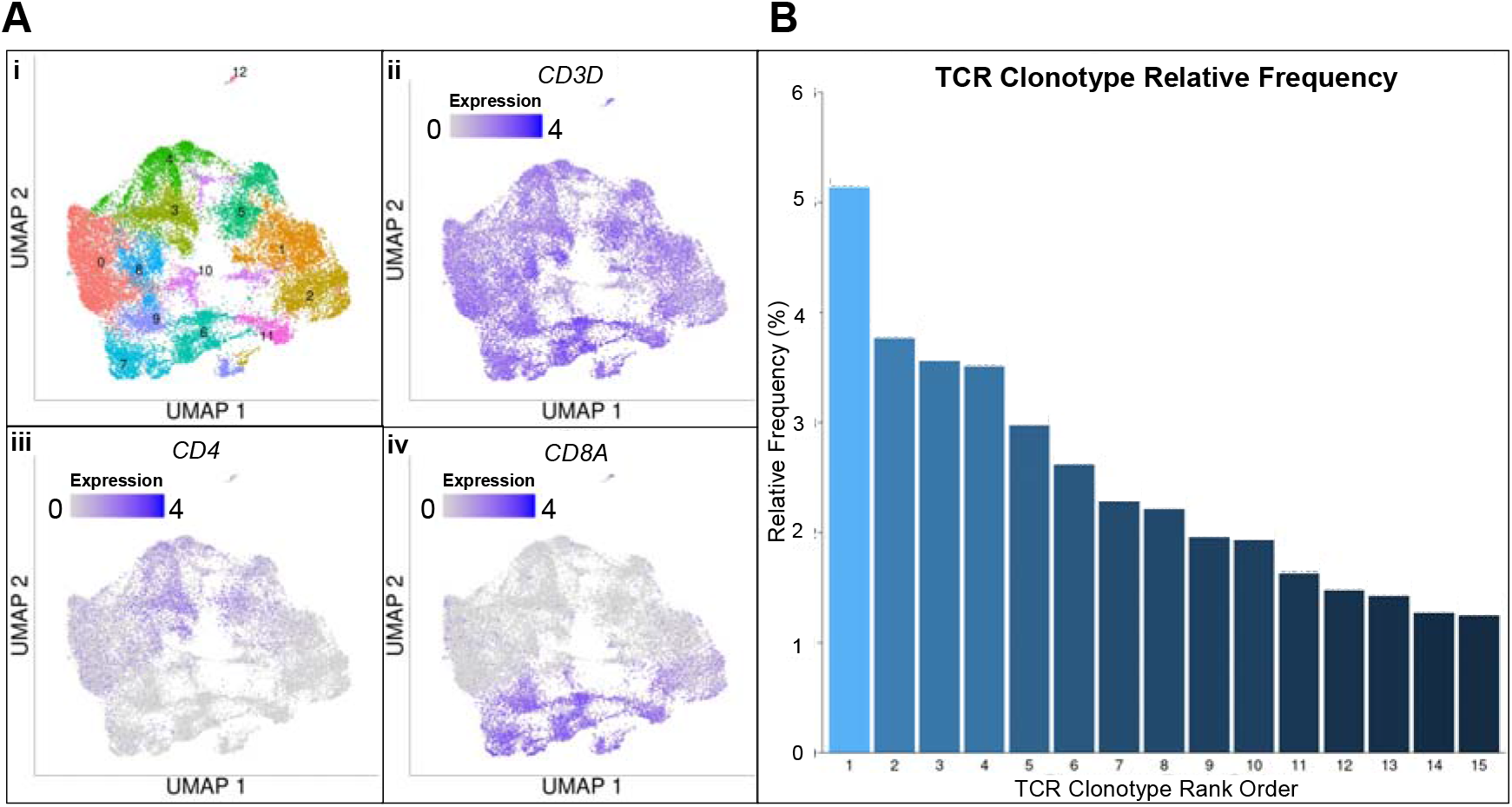
Identified TCRB⍰β clonotypes from CRC patient tumors. UMAP plots of T cells sequenced from colorectal cancer include the **A) i)** total number of unsupervised clusters, **ii)** relative *CD3D* expression, **iii)** relative *CD4* expression, **iv)** relative *CD8A* expression. TCR⍰β clonotypes were identified from the subset of T cells with high CD8 expression and the relative frequency **B)** of those clonotypes. Raw single cell sequencing data from reference (21).

To determine the best method for generating TCR-pMHC starting configurations, we generated the configurations by either Modeller (35,36) or ColabFold (32–34). The structures generated in Modeller utilized the DMF5 TCR bound to the HLA-A2 (MHC) restricted MART1 (PDB:3QDJ) (52) as a template structure, and the ColabFold structures were predicted from trained neural networks (32–34). The resulting starting configurations were then solvated in all-atom molecular dynamics simulations at physiological conditions (see *Methods*) for 150-300 ns to equilibrate the protein structures. Equilibrium is indicated by the flattening of the root mean square deviation (RMSD) from the initial configuration with fluctuations less than 0.2 nm for the entire TCR-pMHC structure (**Figure S1**). A bias-variance trade-off algorithm (45) was used to automate the detection of the equilibrated TCR-pMHC structure (**Figure S1**). The equilibration time required for TCR1 (249 & 9 ns), TCR2 (127 & 61 ns), and TCR3 (149 & 136 ns) from Modeller and ColabFold, respectively, demonstrates ~61% reduction in computational cost — on average — using ColabFold.

We next evaluated the structural similarity of the TCR-pMHC structures throughout the equilibration using the GROMOS clustering algorithm (**Figure S2**) (72). Using a C^⍰^ RMSD cutoff of 0.2 nm, the top ten equilibrated clusters contain most of the configurations for TCR1 (86.8% and 98.3%), TCR2 (92.7% and 97.0%), and TCR3 (71.0% and 97.7%) from Modeller and ColabFold, respectively. The top cluster contains a plurality of structures and occurs after the estimated equilibrium time indicating not a single, but a set of converged TCR-pMHC structures (**Figure S2**).

To evaluate the structural similarity at the TCR-pMHC interface of TCR1, TCR2, and TCR3, we selected and aligned a subset of the equilibrated structures (12 structures — 6 Modeller + 6 ColabFold — see *Methods)* (**Figure 3A-C**). An all-to-all structural alignment (72 unique comparisons) was performed to calculate the pairwise RMSD (after equilibration) within and between TCR-pMHC structures generated by Modeller and ColabFold. The average RMSD for TCRs generated within either Modeller or ColabFold is consistent for TCR1-pMHC (0.19 ± 0.04 nm), TCR2-pMHC (0.20 ± 0.07 nm), and TCR3-pMHC (0.19 ± 0.05 nm). However, there is a consistent increase in average RMSD when comparing equilibrated structures between Modeller and ColabFold for TCR1-pMHC (0.41 ± 0.05 nm), TCR2-pMHC (0.40 ± 0.05 nm), and TCR3-pMHC (0.33 ± 0.04 nm) (**Figure S3-S5**). The increase in RMSD occurs despite selecting TCR-pMHCs from distinct equilibrium clusters (**Figure S2**). The increase in configuration dissimilarity between Modeller and ColabFold at the TCR-pMHC interface can be visualized by the aligned and overlayed structures (**Figure 3A-C**). To investigate the relative fluctuations of the substructures at the TCR-pMHC interface, the root mean square fluctuations were calculated at equilibrium for CDR3⍰, CDR3β, and the CEA peptide (**Figure 3A-C**). Fluctuations for all TCRs and TCR substructures are approximately 0.10 nm.

**Figure 3:**
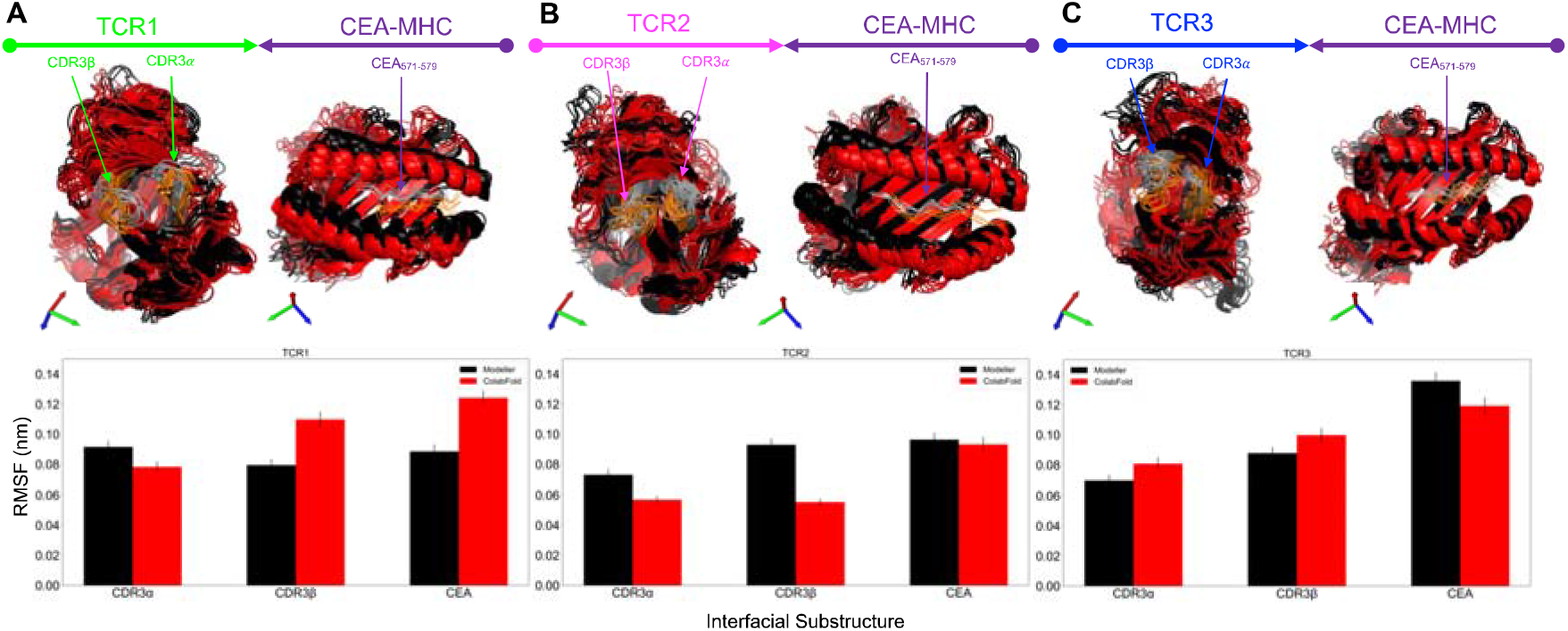
Equilibrated patient-specific TCRs bound to CEA_571-579_ pMHC. The three most frequently found TCRs from 16 CRC patients were predicted (using Modeller or ColabFold) and equilibrated at physiological conditions using molecular dynamics simulations. The most frequent TCRs: TCR1 **(A)**, TCR2 **(B)**, and TCR3 **(C)** are displayed with colors green, magenta, and blue, respectively. The starting structures were created from Modeller (black) and ColabFold (red) and 12 TCR-pMHC structures (6 from Modeller + 6 from ColabFold) are aligned after equilibration with the TCR (on the left, in respective color) and pMHC (on the right, in purple). In addition, the mutated substructures of the TCR and pMHC are indicated by arrows and highlighted in the following colors: CDR3⍰, CDR3β, and CEA_571-579_ (Modeller: gray & ColabFold: orange). After structural equilibration, the root mean square fluctuation (RMSF) for each TCR are calculated for the regions that were mutated: CDR3⍰, CDR3β, and CEA_571-579_ (bottom). The RMSF is calculated for both the Modeller (black) and ColabFold (red) generated starting structures.

After structural equilibration, the number of molecular-level interactions between the TCRs and pMHC were evaluated to assess potential differences between TCRs, and thus provide insight into potential methods to rank the TCRs (**Figure 4**). We selected hydrogen bonds (H-Bonds) and Lennard-Jones contacts (LJ-Contacts) between the TCRs and pMHC to understand the relative importance of coulombic and hydrophobic interactions. The number of hydrogen bonds (**Figure S6**) and the number of Lennard-Jones contacts (**Figure S7**) were calculated as a function of simulation time, and these plots were used to calculate the probability densities. The probability density of interactions is an index to describe the relative likelihood of interactions that occur at any timepoint in equilibrium. Our results demonstrate that TCR2 (expected value: 14.2) is significantly more likely to have more hydrogen bonds than TCR1 (expected value: 12.5) and TCR3 (expected value: 11.7) for structures generated by Modeller (**Figure 4**). In contrast, for structures generated by ColabFold, TCR1 (expected value: 10.5) and TCR3 (expected value: 10.9) are significantly more likely to have more hydrogen bonds than TCR2 (expected value: 6.8) (**Figure 4A**). In addition, TCR2 (expected value: 404) is more likely to have more Lennard-Jones contacts than TCR1 (expected value: 279) and TCR3 (expected value: 294) for structures generated by Modeller. Consistently, for structures generated by ColabFold, TCR2 (expected value: 328) is more likely to have more Lennard-Jones contacts than TCR3 (expected value: 294) and TCR1 (expected value: 306) (**Figure 4B**). Despite TCR1 being detected more frequently in patient tumors (**Figure 2B**), TCR2 may have more binding interactions to CEA571-579 pMHC at equilibrium (**Figure 4C**).

**Figure 4:**
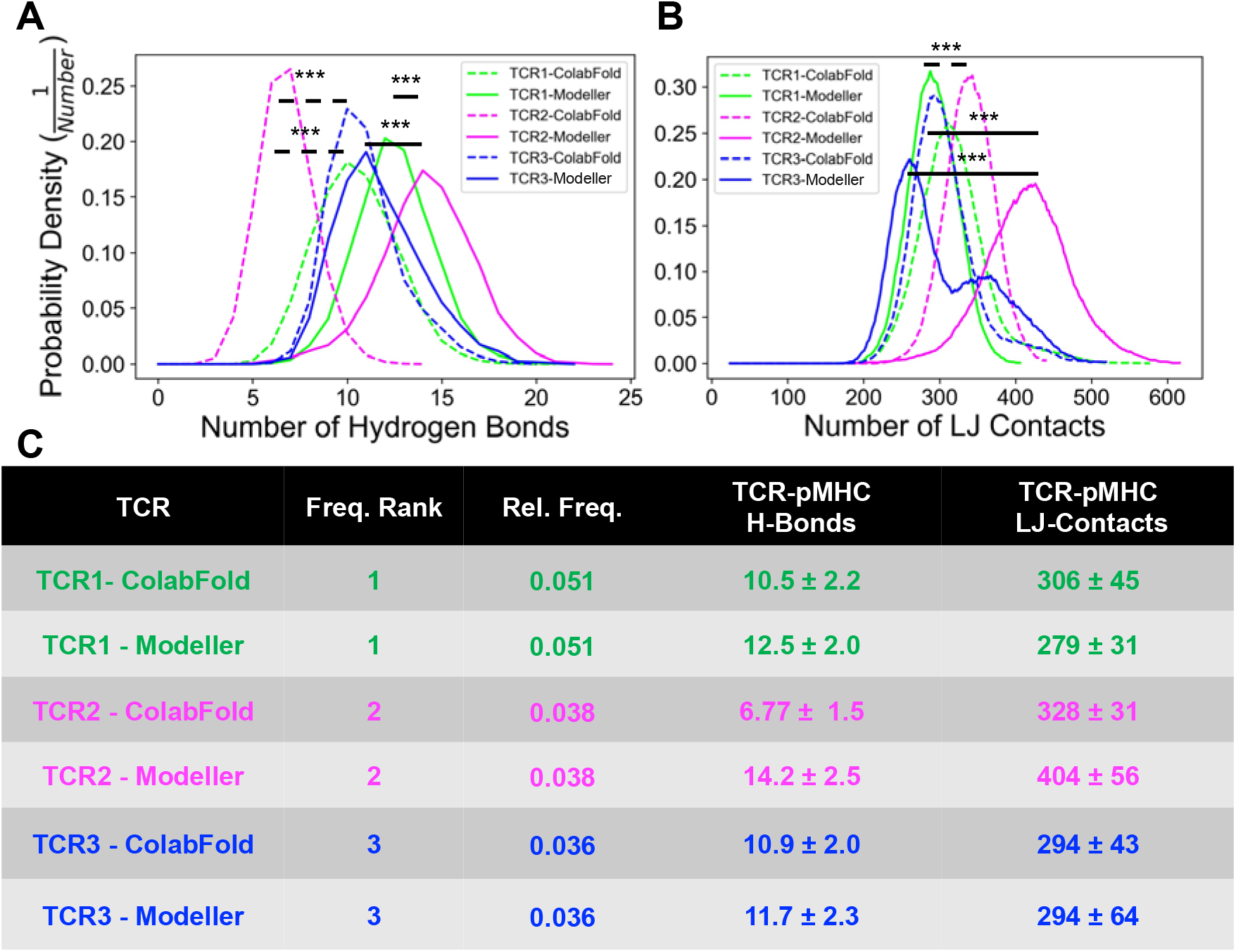
Interactions of patient specific TCRs with pMHC in equilibrium. **(A)** The probability density for the number of hydrogen bonds between the pMHC and TCR1 (green), TCR2 (magenta), and TCR3 (blue), respectively. **(B)** The probability density for the number of Lennard-Jones Contacts between the pMHC and TCR1 (green), TCR2 (magenta), and TCR3 (blue), respectively. The interaction distributions after equilibration are separated for Modeller (solid line) and ColabFold (dashed line). The expected value and standard deviation of H-Bonds and LJ-Contacts is displayed in panel **(C)** with the TCRs ranked by relative frequency found in CRC patients. The number of interactions throughout the simulations were statistically compared: *p<0.05, **p<0.01, ***p<0.001 by one-way ANOVA followed by Tukey-HSD post-hoc test. Statistical significance was only displayed for comparisons with Cohen effect size d>0.5. Significance was displayed by solid and dashed lines for Modeller and ColabFold comparisons, respectively.

## DISCUSSION

Using a combination of single cell sequencing of T cells derived from patient tumors, protein structure prediction algorithms, and MD simulations, we present a pipeline to rank TCRs based on their molecular level interactions with a target pMHC at equilibrium. To commence a pipeline to assess TCRs *in silico*, we chose only the three most frequently expressed clonotypes as a case study. Although, the selection of the most frequent TCRs from the T cell repertoire in the tumor microenvironment is somewhat arbitrary, the relatively expanded T cells in the tumor microenvironment are more likely to be immunogenic to the tumor than a random selection of low frequency clones. Nonetheless, our pipeline is easily adapted to selecting clonotypes from alternate sources (e.g., peripheral blood), or alternate strategies (e.g., selecting from the entire clonal pool in the tumor microenvironment).

Our study also presented an opportunity to assess two fundamentally different protein-protein structure prediction tools: ColabFold — a recently released trained deep learning network based on AlphaFold(32,33); and Modeller — a traditional template-based prediction tool. We found that the MD equilibration of 3D atomic coordinates of TCR-pMHC structures was ~ 2.5X faster using ColabFold generated structures (**Figure S1**). This finding is based on a small number of protein-protein structures, and there was significant variation. Nonetheless, our results demonstrate that ColabFold may be a superior computational tool for protein-protein structure prediction, and thus may have implications on the scale-up of assessing a larger set (i.e., thousands) of TCRs generated from sequencing data.

To automate and remove human bias from determining the required simulation time to reach an equilibrated TCR-pMHC structure, we used a variance-bias trade-off algorithm(45) and required a minimum time of 50 nanoseconds in equilibrium (**Figure S1**). In addition, we performed a cluster analysis to identify the set of converged clusters after equilibration (**Figure S2**). Moreover, we found that the rootmean-square fluctuations after equilibration for CDR3⍰, CDR3β, and the CEA peptide (**Figure 3A-C**) were approximately 0.10 nm. The fluctuations for CDR3⍰, CDR3β, and peptide are consistent with equilibrated TCRs with a known crystal structure (28), and thus indicative of equilibrium.

After equilibration, we assessed several clusters of configurations and found that within a protein-protein structure predictor (i.e., Modeller or ColabFold) there is a pairwise RMSD of ~0.20 nm. Interestingly, across structure predictors there is an increase in pairwise RMSD ~0.40 nm (**Figure S3-S5**). This trend is consistent for TCR1, TCR2, and TCR3 indicating that the structure prediction method can influence the set of equilibrated TCR-pMHC configurations, and molecular level interactions. We found that TCR2 had more hydrogen bonds compared to TCR1 and TCR3 when generated by Modeller, but less hydrogen bonds when generated by ColabFold (**Figure 4A**). These results indicate that the differences in configurations generated by Modeller and ColabFold can also influence the number of molecular level interactions between the TCR and pMHC. We did observe consistent results between Modeller and ColabFold for the number Lennard-Jones contacts across the three TCRs (**Figure 4B**). The probability density of hydrogen bonds and Lennard-Jones contacts may provide a rudimentary criterion to rank TCRs based on their relative strength of interaction (28) at equilibrium. For example, TCR2 may be a more ideal target to the CEA571-579 pMHC because of the consistent increase in Lennard-Jones contacts. To develop a refined TCR ranking index, future work will require a comprehensive dataset that will assess the physiochemical properties of TCR-pMHC interactions that determine immunogenicity.

### Conclusions

The identification of tumor-specific TCRs will be augmented by computational methodologies that accurately rank TCRs based on the immunogenic response to a target pMHC. We have integrated next-generation sequencing with protein-protein structure prediction and MD to introduce a potential pipeline to evaluate TCRs. We found that ColabFold outperforms Modeller (~2.5X) in the required simulation time to generate equilibrated TCR-pMHC structures, and thus may be a superior computational tool to utilize in a pipeline built to assess and predict TCR immunogenicity. In addition, the protein-protein structure prediction method influences the set of equilibrated configurations and the number of interactions between the TCR and pMHC, and thus may impact the accuracy of predicting TCR-pMHC bond strength or immunogenic response.

## AUTHOR CONTRIBUTIONS

ZAR performed the simulations, analyzed, and interpreted the data, and wrote the manuscript. MBC analyzed and interpreted the RNA sequencing data and wrote the manuscript. RF designed the experiments, analyzed and interpreted the data, wrote the manuscript, and secured computer time. SCG designed the experiments, analyzed, and interpreted the data, wrote the manuscript, and secured the funding.

## ACKNOWLDGEMENTS

Single cell V(D)J RNA sequencing data was a generous gift from Professor Arnold Han at Columbia University. Simulations were performed on the hpc1/hpc2 clusters at UC Davis. This work was supported in part by startup funding to SCG form the Department of Biomedical Engineering.

## DATA AVAILABILITY

The single cell RNA-Seq data used in this study has previously been made available at ArrayExpress #E-MTAB-9455. TCR-pMHC structures generated from proteinprotein structure predictors have been made available in a Dryad repository: https://doi.org/10.25338/B83S70. In addition, the structures, box sizes, and atom counts are all deposited in this repository. All scripts relevant to the production of figures have been made available on a Github repository: https://github.com/zrollins/TCR_homology.git.

