## Supporting Information for "Automated protein-protein structure prediction of the T cell receptor-peptide major histocompatibility complex"

**This PDF includes:**

**Supplementary Figures**

Fig. S1. Equilibration of the TCR-pMHC systems.
Fig. S2. Cluster analysis of the TCR-pMHC systems.
Fig. S3. TCR1 pairwise RMSDs for clustered Modeller and ColabFold structures.
Fig. S4. TCR2 pairwise RMSDs for clustered Modeller and ColabFold structures.
Fig. S5. TCR3 pairwise RMSDs for clustered Modeller and ColabFold structures.
Fig. S6. Hydrogen bonds for the TCR-pMHC systems.
Fig. S7. Lennard-Jones contacts for the TCR-pMHC systems.

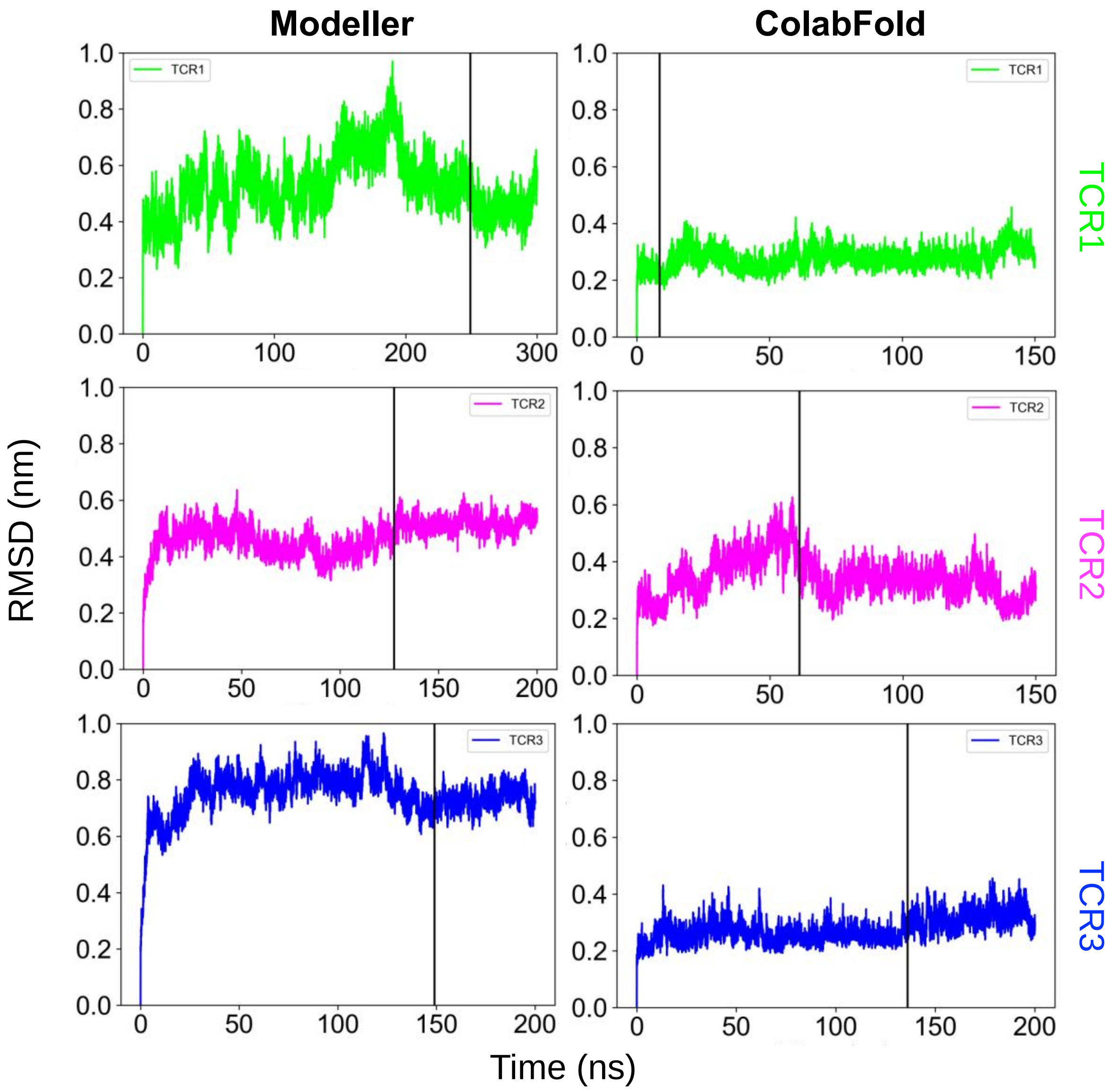


**Figure S1: Equilibration of the TCR-pMHC systems.** The root mean square deviation (RMSD) is calculated from the initial configuration and plotted versus simulation time (x-axis). RMSD is an all-atom calculation on the entire TCR-pMHC structure. The initial configurations generated by protein-protein structure predictors are identified by columns: Modeller (left) and ColabFold (right). The TCRs are identified by rows: TCR1 (top, green), TCR2 (middle, magenta), and TCR3 (bottom, blue). The equilibrated time-point determined by the variance-bias trade-off algorithm is displayed by the black vertical line in each panel.

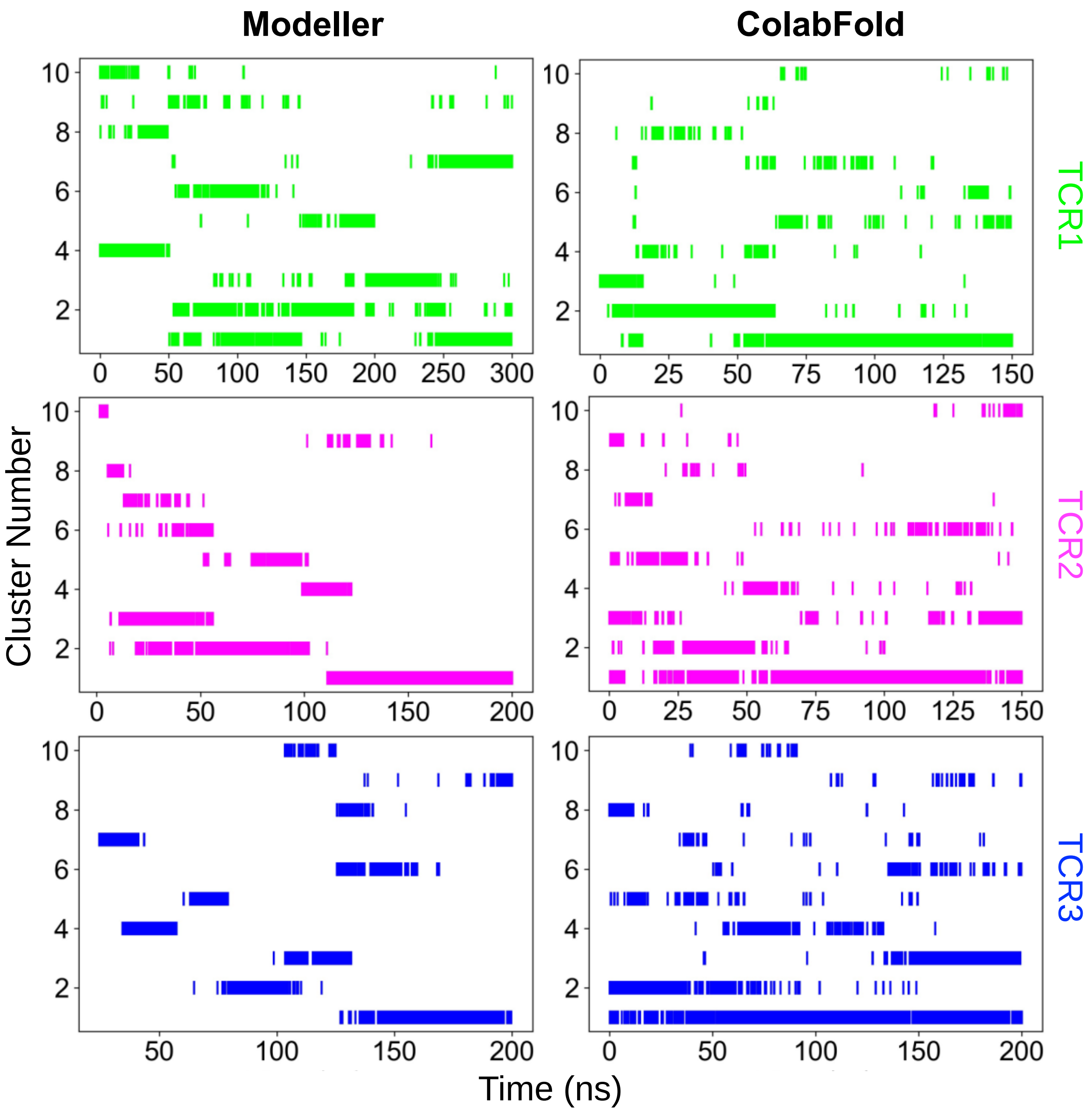


**Figure S2: Cluster analysis of the TCR-pMHC systems.** The GROMOS clustering algorithm with a $C^{\alpha}$ RMSD cutoff of 0.20 nm was used to identify dominant structural configurations during equilibration. The cluster numbers (y-axis) are ordered by the number of configurations in each cluster (i.e., one is the most dominant) and plotted versus simulation time (x-axis). From Modeller, 70, 45, and 102 clusters were identified with 86.8% (26030/30000), 92.7% (18540/20000), and 71.0% (14206/20000) in the top ten clusters and 26.6% (7980/30000), 38.8% (7763/20000), and 23.1% (4621/20000) in the top cluster for TCR1, TCR2, and TCR3, respectively (**left**). From ColabFold, 25, 28, and 31 clusters were identified with 98.3% (14749/15000), 97.0% (14550/15000), and 97.7% (19530/20000) in the top ten clusters and 54.4% (8158/15000), 55.1% (8263/15000), and 50.7% (10133/20000) in the top cluster for TCR1, TCR2, and TCR3, respectively (**right**). The TCRs are identified by rows: TCR1 (top, green), TCR2 (middle, magenta), and TCR3 (bottom, blue).

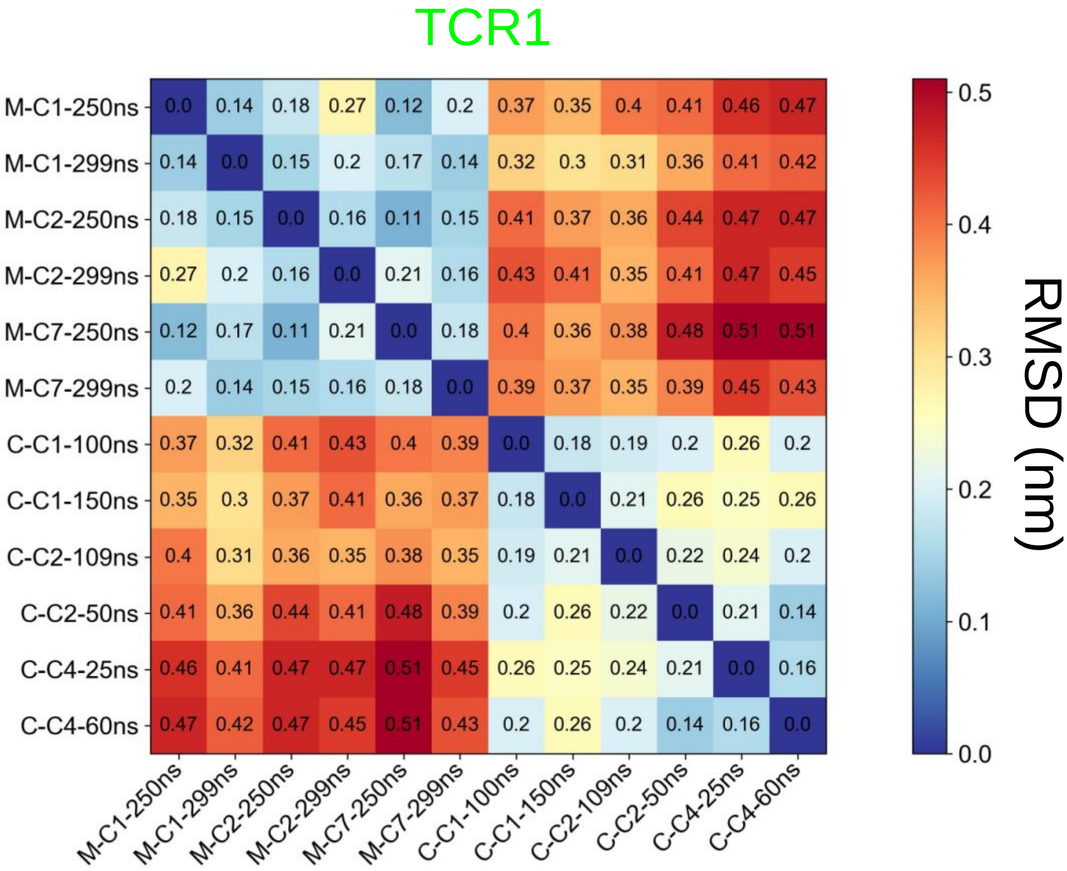


**Figure S3: TCR1 pairwise RMSDs for clustered Modeller and ColabFold structures.** The selection of 12 TCR-pMHC structures (6 Modeller + 6 ColabFold) were chosen to be from distinct clusters after equilibration. The pairwise RMSD between selected structures is displayed with the following format: protein-protein structure predictor (i.e., M=Modeller or C=ColabFold) — cluster number from Figure S2 (e.g., C1= cluster 1) — simulation time (e.g.,100 ns). The RMSD is calculated on the $C^{\alpha}$ atoms and scale bar is displayed in nanometers (**right**). The clusters were selected because they were after the simulation time required for equilibrium (e.g., for Modeller TCR1, cluster 1,2, & 7 were chosen because clusters 3-6 are only dominant before reaching equilibrium at 249 ns simulation time).

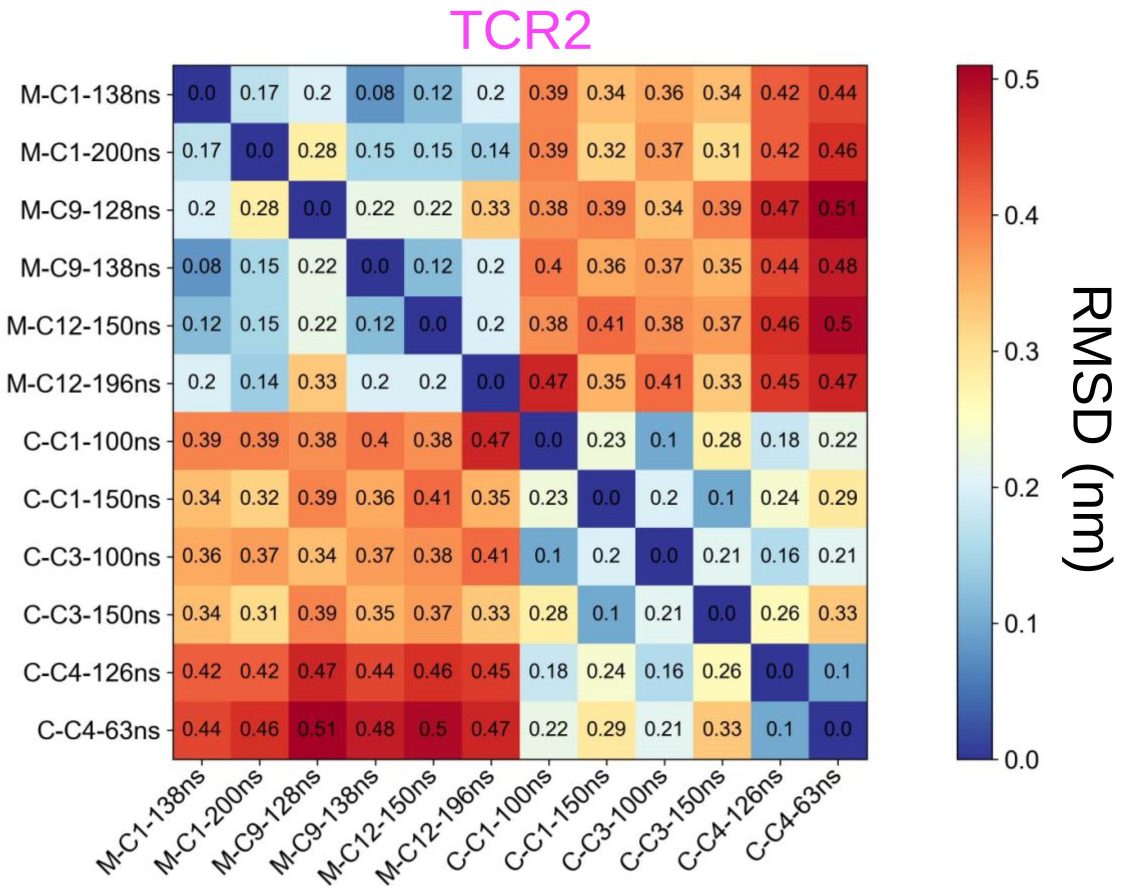


**Figure S4: TCR2 pairwise RMSDs for clustered Modeller and ColabFold structures.** The selection of 12 TCR-pMHC structures (6 Modeller + 6 ColabFold) were chosen to be from distinct clusters after equilibration. The pairwise RMSD between selected structures is displayed with the following format: protein-protein structure predictor (i.e., M=Modeller or C=ColabFold) — cluster number (e.g., C1= cluster 1) — simulation time (e.g.,100 ns). The RMSD is calculated on the $C^{\alpha}$ atoms and scale bar is displayed in nanometers (**right**). The clusters were selected because they were after the simulation time required for equilibrium (e.g., for Modeller TCR2, cluster 1,9, & 12 were chosen because clusters 2-8 &10-11 are only dominant before reaching equilibrium at 127 ns simulation time).

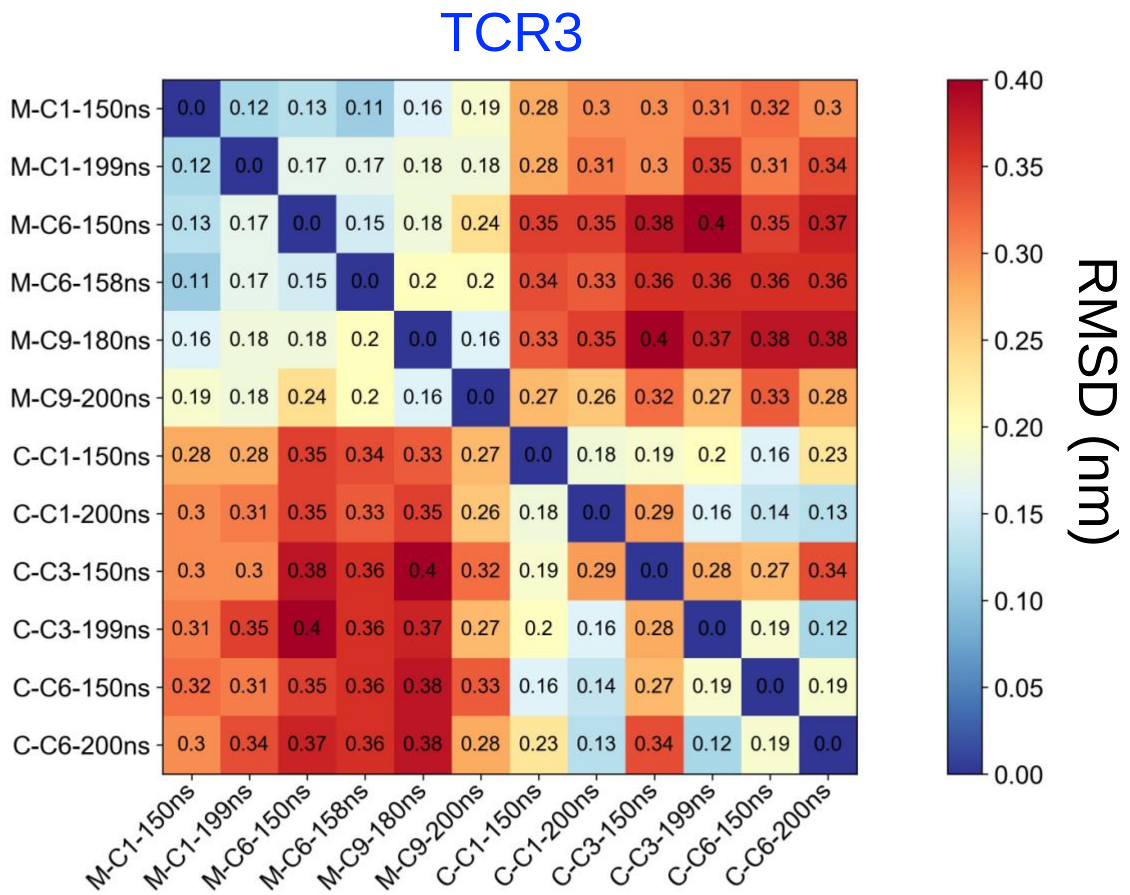


**Figure S5: TCR3 pairwise RMSDs for clustered Modeller and ColabFold structures.** The selection of 12 TCR-pMHC structures (6 Modeller + 6 ColabFold) were chosen to be from distinct clusters after equilibration. The pairwise RMSD between selected structures is displayed with the following format: protein-protein structure predictor (i.e., M=Modeller or C=ColabFold) — cluster number (e.g., C1= cluster 1) — simulation time (e.g.,100 ns). The RMSD is calculated on the $C^{\alpha}$ atoms and scale bar is displayed in nanometers (**right**). The clusters were selected because they were after the simulation time required for equilibrium (e.g., for Modeller TCR3, cluster 1,6, & 9 were chosen because clusters 2-5 & 7-8 are only dominant before reaching equilibrium at 149 ns simulation time).

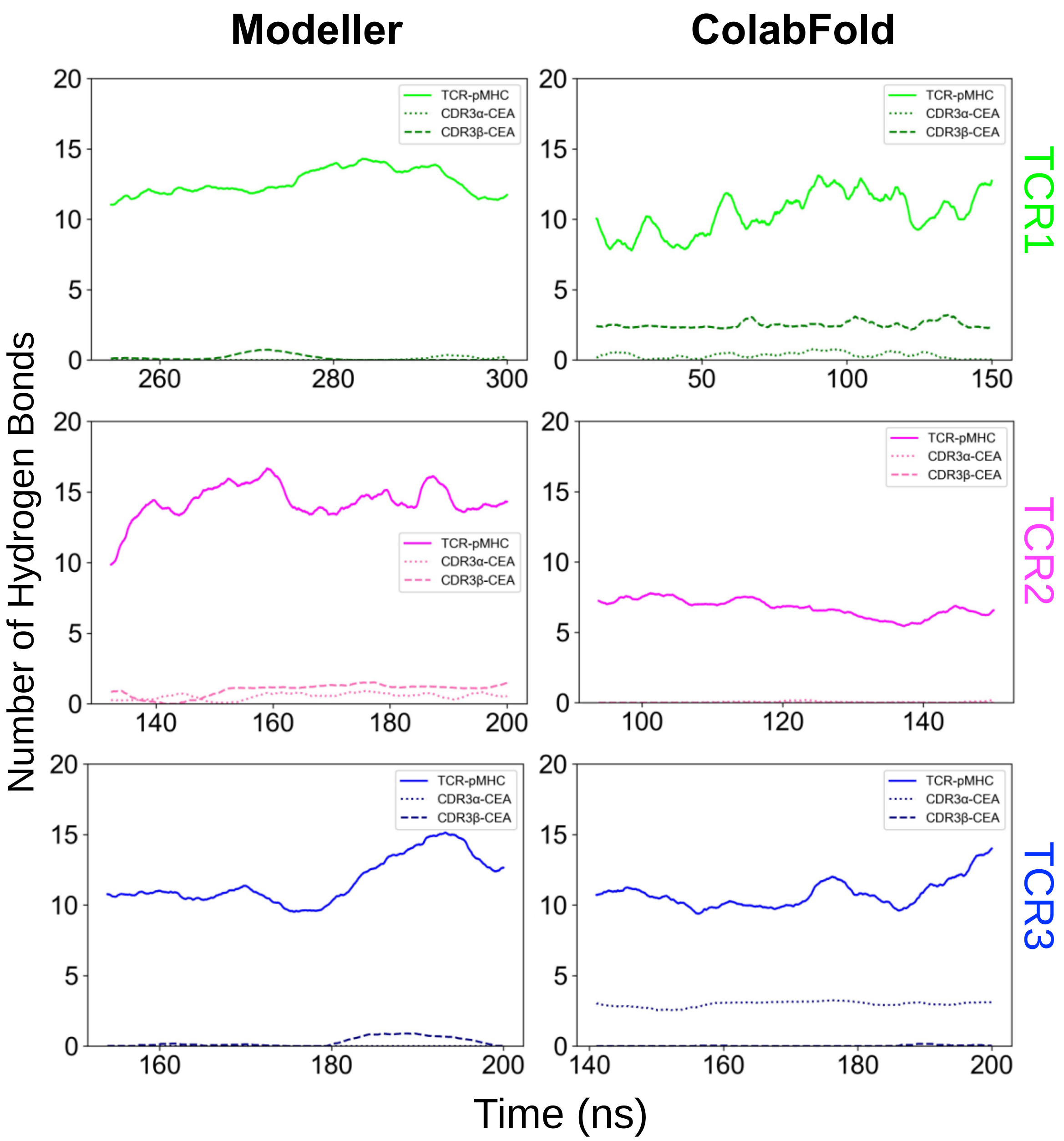


**Figure S6: Hydrogen bonds for the TCR-pMHC systems.** The number of hydrogen bonds (y-axis) is plotted versus simulation time (x-axis) after the determined equilibrated time-point. The initial configurations generated by protein-protein structure predictors are identified by columns: Modeller (left) and ColabFold (right). The TCRs are identified by rows: TCR1 (top, green), TCR2 (middle, magenta), and TCR3 (bottom, blue). The number of hydrogen bonds between the TCR-pMHC (solid line), CDR3α-CEA (dotted line), and CDR3β-CEA (dashed line) are plotted versus simulation time.

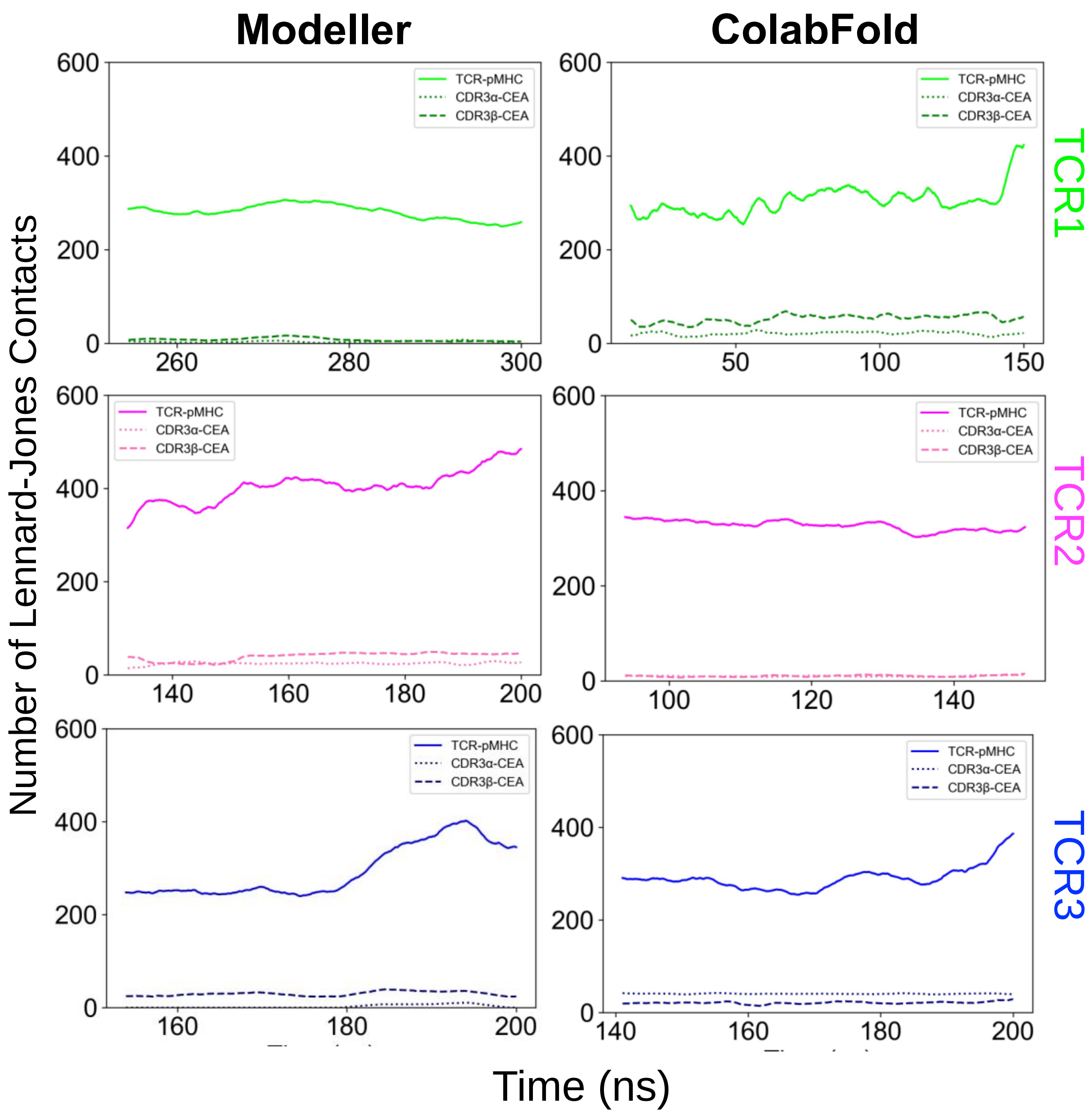


**Figure S7: Lennard-Jones contacts for the TCR-pMHC systems.** The number of Lennard-Jones contacts (y-axis) is plotted versus simulation time (x-axis) after the determined equilibrated time-point. The initial configurations generated by protein-protein structure predictors are identified by columns: Modeller (left) and ColabFold (right). The TCRs are identified by rows: TCR1 (top, green), TCR2 (middle, magenta), and TCR3 (bottom, blue). The number of Lennard-Jones contacts between the TCR-pMHC (solid line), CDR3α-CEA (dotted line), and CDR3β-CEA (dashed line) are plotted versus simulation time.
